# Regulatory Landscape Enrichment Analysis (RLEA) using gaiaAssociation

**DOI:** 10.1101/2023.10.11.561933

**Authors:** Eric A. Sosa, Samuel Rosean, Dónal O’Shea, Srilakshmi M. Raj, Cathal Seoighe, John M. Greally

**Author notes:** Equal Contribution (Eric A. Sosa, Samuel Rosean, Dónal O’Shea).

## Abstract

**Motivation:** To understand whether sets of genomic loci are enriched at the regulatory loci of one or more cell types, we developed the gaiaAssociation package to perform Regulatory Landscape Enrichment Analysis (RLEA). RLEA is a novel analytical process that tests for enrichment of sets of loci in cell type-specific open chromatin regions (OCRs) in the genome.

**Results:** We demonstrate that the application of RLEA to genome-wide association study (GWAS) data reveals cell types likely to be mediating the phenotype studied, and clusters OCRs based on their shared regulatory profiles. GaiaAssociation is Python code that is freely available for use in functional genomics studies.

**Availability and Implementation:** Gaia Association is available on PyPi (https://pypi.org/project/gaiaAssociation/0.6.0/#description) for pip download and use on the command line or as an inline Python package. Gaia Association can also be installed from GitHub at https://github.com/GreallyLab/gaiaAssociation.

## Introduction

Despite the immense number of genome-wide association studies (GWAS) performed to study human diseases and traits, prioritizing the resulting variants for functional characterization remains a difficult task. Most DNA sequence variants implicated in GWAS occur in the non-coding majority of the human genome, and have been explored with functional genomic approaches to identify those variants with effects on gene expression or transcriptional regulatory processes (Uffelmann *et al*., 2021). These variants with gene regulatory function are often considered candidates for mediating the organismal phenotype.

Disease-relevant functional variants are likely to be more frequent in regulatory regions that are active in the cell or tissue types mediating the condition. With the increasing availability of data from chromatin state assays such as ATAC-seq (Buenrostro *et al*., 2013) that enable the identification of loci involved in cell type-specific gene regulation, we hypothesized that GWAS results could be used to identify disease-mediating cell types (Maurano *et al*., 2012; Soskic *et al*., 2019; Hauberg *et al*., 2020).

To test for enrichment of sets of loci in the regulatory elements of different cell types, we developed the gaiaAssociation package to perform Regulatory Landscape Enrichment Analysis (RLEA). The implementation of RLEA partitions the genome into equally sized windows and then models the number of overlaps between single nucleotide variants (SNVs) or small indels and ATAC-seq peaks in each window as a binomial random variable. For the entire genome, the number of overlaps is modeled as a sum of independent binomial random variables, which can be estimated using saddlepoint approximation (Liu *et al*., 2017). This method reveals candidate cell types mediating the effects of genetic variants on the phenotype, a valuable post-GWAS insight, while clustering cell types by their OCR enrichment profiles.

We apply this RLEA approach to data from the NHGRI-EBI GWAS catalog (Sollis *et al*., 2023), comparing enrichments across the regulatory landscapes of 44 cell types. We show how RLEA yields insights into the cell types potentially mediating the traits and diseases studied by GWAS.

## Methods

### 2.1 Genomic resources

We applied RLEA to identify the cell types through which causal variants impact a trait or disease. We used ATAC-seq data from 44 healthy cell types, derived from the ATACdb database (Wang *et al*., 2021) and from a published dataset of human brain samples (Hauberg *et al*., 2020). The NHGRI-EBI GWAS catalog was used to identify 303 studies with ≥200 loci reaching genome-wide significance, from which we selected 18 (**Table S1**), choosing a variety of traits and diseases that could potentially be mediated by one or more of the 44 cell types for which we had ATAC-seq data. All ATACdb datasets were converted to the current GRCh38.p13 genome build using the UCSC Genome Browser *LiftOver* function.

### 2.2 Software interface for data input

The gaiaAssociation package requires four inputs for analysis: 1) a folder containing the chromatin regulatory region data of interest in bed text (.txt) format, 2) a folder containing the genomic loci of interest in tab-separated values (.tsv) or comma-separated values (.csv) format, 3) a .csv file containing the chromosome sizes of the desired genome build, and 4) a folder for the output of gaiaAssociation analyses.

### 2.3 Data pre-processing

The most straightforward example of the use of gaiaAssociation is represented by the input of a list of loci to test for overlap with OCRs from multiple cell types. The package supports SNVs and small indels, not larger copy number variants, with the primary goal of supporting data typical of GWAS results. When entering multiple sets of genomic loci, the user can define cutoff values to filter for a minimum size requirement for a dataset, for example. We used a minimum cutoff of 200 genomic loci as a conservative threshold.

GaiaAssociation also permits the user to incorporate their own ATAC-seq loci or regulatory loci defined by other genome-wide assays. The software tests the relatedness of the loaded ATAC-seq profiles as a pre-processing step. The relationship between each pair of cell types is defined by calculating a weight, using the following formula:

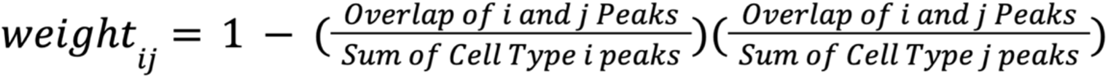

This weight matrix can be used to generate a dendrogram, based on the Euclidean distance metric and hierarchical clustering, using the SciPy8 algorithm (Virtanen *et al*., 2020), highlighting the similarities in chromatin organization between cell types.

GaiaAssociation also allows prioritization of regulatory regions that are cell-type specific. The degree of OCR cell type specificity can be generated from the “peak uniqueness value” function. This function removes overlapping OCRs between cell types, allowing filtering prior to analysis as an option. If this function is provided a uniqueness cutoff value of 3, for example, then only OCRs found in three cell types or fewer are considered, removing OCRs shared by four or more cell types.

### 2.4 Regulatory Landscape Enrichment Analysis (RLEA)

To calculate the sum of independent but non-identical binomial random variables, RLEA takes the user-provided chromosome size file and divides the chromosomes into windows of equal length, as close to the user-defined window size as possible. For each window in which there is at least one query loci /SNP, the number of overlaps between the query set and the subject regions in the absence of enrichment is modeled as a binomial random variable. This binomial random variable is determined by OCR overlap percentage and loci count. Thus, for the entire genome, the number of overlaps is modeled as a sum of independent binomial random variables. This distribution is estimated using saddlepoint approximation (Liu *et al*., 2017), allowing the query set to be tested for enrichment, given the observed number of overlaps. For each window, the proportion that is occupied by OCRs is calculated for each cell type, and the total number of loci within that window is found. In the case of loci that overlap the OCR and have a length greater than one base pair, the size of each OCR in the window is extended to include the length of these extended loci. This is a slight adjustment to the more conventional single base pair loci calculation which is made to account for the possibility that an indel could land outside the OCR region but still overlap with it. This proportion of OCR coverage and the count of loci that overlap OCRs within that window, along with the total number of loci that overlap OCRs genome-wide, are used to calculate a p-value for global enrichment using the sinib package (Liu *et al*., 2017), which has been translated from R to Python and included with gaiaAssociation. This p-value is calculated for each cell type (*n* values) and loci dataset (*m* values) combination, to create an *n* x *m* matrix which describes the relative enrichment of these loci in each cell type.

## Results

### RLEA of GWAS loci

RLEA was used to analyze a subset of traits for which GWAS have generated at least 200 loci reaching genome-wide significance (**Figure 1**). We chose 18 traits or diseases that could be mediated by the 44 selected cell types, from which we highlighted nine associations in Figure 1. These examples demonstrate how RLEA links GWAS results with cell types potentially mediating the studied trait or disease. Some associations appear to support roles for specific cell types, such as the association of type 2 diabetes mellitus genetic risk loci with pancreatic islets, autoimmune and inflammatory disorders with immune cells, counts of different circulating blood cell types with regulatory regions of those cells, and some brain cell types with psychiatric or neurodevelopmental phenotypes. Some of these associations are consistent with prior studies linking GWAS variants with putative immune cell regulatory loci in autoimmune or inflammatory diseases (Maurano *et al*., 2012; Soskic *et al*., 2019) or psychiatric disease and neurodivergent traits (Dorph-Petersen *et al*., 2007; Galvez-Contreras *et al*., 2020).

**Figure 1.**
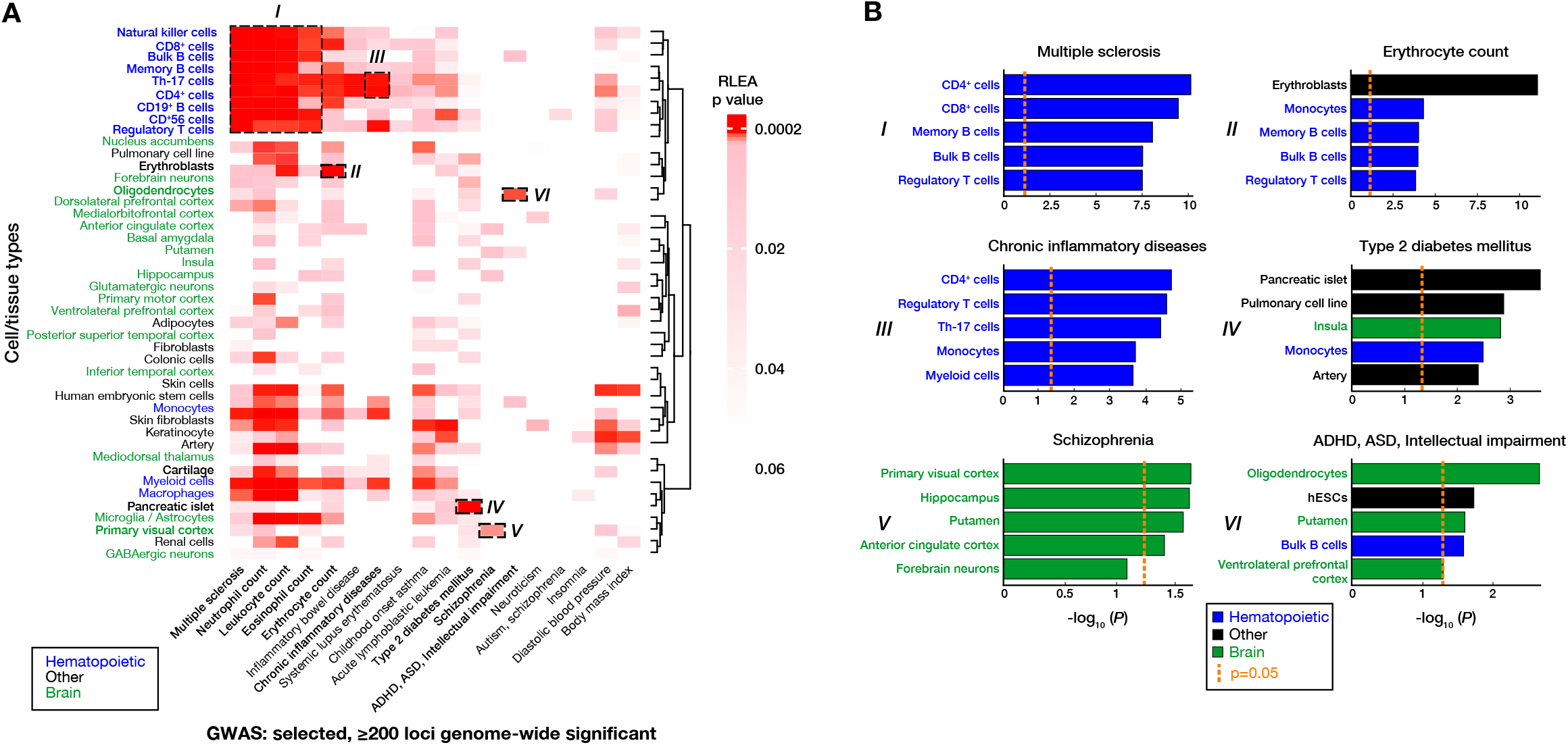
Regulatory Landscape Enrichment Analysis (RLEA) allows the association of sets of genomic loci with regulatory loci of different cell types. We show the results of enrichment of significant loci from 18 GWAS and the *cis*-regulatory regions of 44 cell types. **(A)** RLEA p values are visualized using a heatmap, using darker red to show the cell types that survive correction for multiple comparisons. We highlight a few associations in particular, (I) the association of multiple sclerosis and blood cell counts with the regulatory loci of a number of immune cell types, (II) erythroblasts with erythrocyte count, (III) Th-17 and CD4+ lymphocytes with chronic inflammatory diseases (ankylosing spondylitis, ulcerative colitis, Crohn’s disease, psoriasis, sclerosing cholangitis), (IV) pancreatic islets with type 2 diabetes mellitus, (V) primary visual cortex with schizophrenia, and (VI) oligodendrocytes with attention deficit hyperactivity disorder (ADHD), autism spectrum disorder (ASD) and intellectual impairment. **(B)** The p values for the top 5 cell type associations for each of the 6 highlighted examples from panel (A) are shown. The orange dashed lines at −log_10_ (*P*) = 1.30 represents a p value cutoff of < 0.05. These analyses illustrate some of the potential applications of RLEA.

Understanding the distribution of associated variants across the *cis*-regulatory regions of different cell types provides valuable clues to the cell type(s) through which they affect a trait or disease, and a clue to the identity of the regulatory elements that should be the focus of studies of non-coding sequence variation. We have provided a novel and efficient analytical resource for RLEA analysis that is freely-available and a useful part of the toolkit for understanding non-coding variants in human traits and diseases.

## Supplementary information

A supplementary table summarizing the datasets analyzed in the present study has been provided and is available at Bioinformatics online.

## Acknowledgements

We thank David Yang from our laboratory for his help in the initial experimental validation of this project.

## Disclosure of Potential Conflicts of Interest

No potential conflicts of interest were disclosed.

**Supplementary Figure 1:**
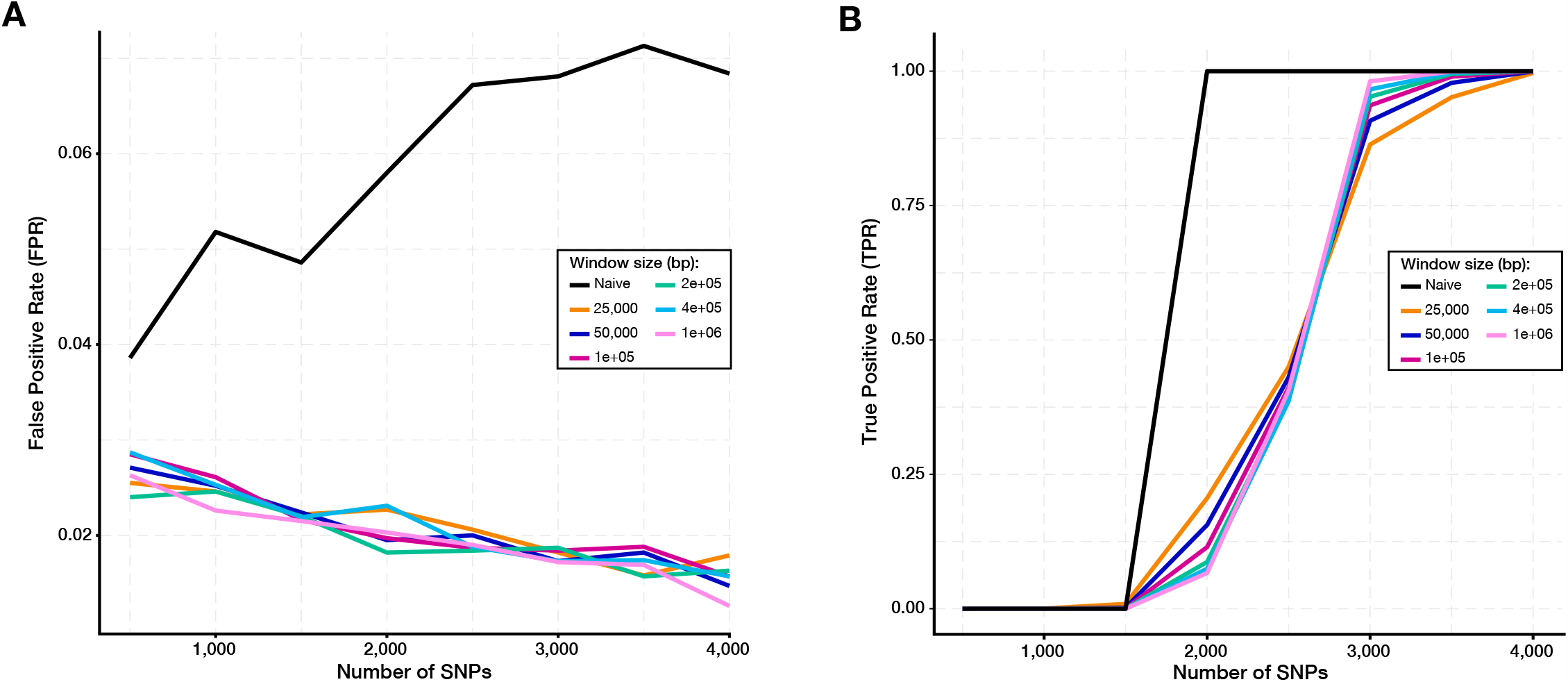
100,000 random simulations of RLEA demonstrate that increased SNPs is inversely related to false positive rate (FPR) and positively corelated to true positive rate (TPR). **(A)** FPR declines as the number of SNPs increases and window size is defined. RLEA FPR was calculated for varying window sizes ranging from 25,000-1,000,000 base pairs (bp) in length. The naïve window size (black line) represents the whole genome in the absence of window selection. **(B)** Increased SNP number and window size are directly correlated with TPR in RLEA. The naïve window TPR plateaus before all defined windows since decreased window sizes are associated with increasingly conservative enrichment. All SNPs in this simulation were selected at random and tested for enrichment in real ATAC-seq peaks.

